# STING polymer structure reveals mechanisms for activation, hyperactivation, and inhibition

**DOI:** 10.1101/552166

**Authors:** Sabrina L. Ergun, Daniel Ferdandez, Thomas M. Weiss, Lingyin Li

## Abstract

How the central innate immune protein, STING, is activated by its ligands remains unknown. Here, using structural biology and biochemistry, we report that the metazoan second messenger 2’3’-cGAMP induces closing of the human STING homodimer and release of the STING C-terminal tail, which exposes a polymerization interface on the STING dimer and leads to the formation of disulfide-linked polymers via cysteine residue 148. Disease-causing hyperactive STING mutations either flank C148 and depend on disulfide formation or reside in the C-terminal tail binding site and cause constitutive C-terminal tail release and polymerization. Finally, bacterial cyclic-di-GMP induces an alternative active STING conformation, activates STING in a cooperative manner, and acts as a partial antagonist of 2’3’-cGAMP signaling. Our insights explain the tight control of STING signaling given varying background activation signals and provide a novel therapeutic hypothesis for autoimmune syndrome treatment.

## Main Text

The stimulator of interferon genes (STING) pathway senses cytosolic double-stranded DNA (dsDNA) which can be a sign of viral or bacterial infection, damaged cells, or erroneous chromosomal segregation of cancerous cells. Upon sensing of dsDNA, the enzyme cyclic-GMP-AMP-synthase (cGAS), cyclizes GTP and ATP to produce the second messenger 2’3’-cyclic-GMP-AMP (cGAMP) (*1-6).* cGAMP binds to and activates the endoplasmic reticulum transmembrane receptor STING, which consists a cytosolic cGAMP binding domain and a four-pass transmembrane domain (*6-10*). Activated STING then serves as an adaptor for kinase TBK1 and transcription factor IRF-3 and leads to IRF-3 phosphorylation and dimerization (*11, 12*). Phosphorylated IRF3 dimers translocate to the nucleus and induce the production and secretion of type I IFNs, which are potent anti-viral, anti-bacterial, and anti-cancer cytokines (*3-5,13, 14).*

STING was originally characterized for its central roles in anti-viral immunity. STING deficient mice are more susceptible to DNA viruses (*13, 16*) and retroviruses including HIV (*13*). STING is now also recognized as a promising target for cancer immunotherapy. Intra-tumoral injection of STING agonists (*17*) exerted remarkable curative effects in multiple synergistic mouse tumor models (*18-22*) and two have since entered clinical trials (Trial IDs NCT03172936 and NCT03010176). Importantly, homozygous loss-of-function STING mutations have not been reported in the human population suggesting that the pathway is essential for survival (http://exac.broadinstitute.org/gene/ENSG00000184584).

Conversely, high levels of STING activation have been implicated in many debilitating autoimmune syndromes such as systemic lupus erythematosus, multiple sclerosis, and Aicardi-Goutières syndrome (*23-26*). In addition, STING pathway hyperactivity is responsible for acute inflammation in myocardial infarction and for chronic inflammation in liver drug toxicity, liver disease, and pancreatitis (*27, 28).* Moreover, six point mutations in STING have been reported in children that cause STING hyperactivity and lead to the autoimmune syndrome STING Associated Vasculopathy with Onset in Infancy (SAVI) (*29-32*). The mechanism of STING activation by its ligands, and how STING is able to balance its essential response to foreign and cancer-derived dsDNA, but not induce autoimmunity, remain major unsolved questions in the field.

To begin to understand how human STING is activated by cGAMP, we sought to use a chemical biology approach and formally investigate whether other small molecule STING binders exert the same activity. In addition to the metazoan cyclic dinucleotide (CDN) cGAMP, other CDNs cyclic-di-GMP (CDG) and cyclic-di-AMP (CDA), which are ubiquitous bacterial signaling molecules, activate the STING pathway in mice (*8, 33, 34*). Mice harboring the null I199N STING mutation (*goldenticket*) have an increased susceptibility to intracellular bacterial pathogens *Listeria monocytogenes* and *Mycobacterium tuberculosis* which produce CDG and CDA respectively (*35-38)*. The roles of CDG and CDA in human STING activation are largely unexplored. Since we and others previously reported that human and mouse STING have drastically different ligand selectivity (*39, 40*), we cannot assume that CDG and CDA are human STING agonists. For example, we previously reported that CDG binds to human STING with ∼130-fold lower affinity than to mouse STING (*39*). Crystallographic studies also showed that cGAMP, but not CDG, induces closing of the human STING dimer to a similar angle as that of mouse STING (*5, 42*). Unlike CDG, CDA has been much less studied in the context of human STING. Although it has been reported that addition of CDA to 293T cells expressing human STING led to STING dependent gene expression, direct binding of CDA and human STING has never been demonstrated.

## Results

### Human STING forms closed dimer angle when bound to cGAMP and CDA

Similar to previously published results (*18*), we found that CDA activates IFN-β production in primary human lymphocytes expressing wildtype (WT) human STING and the 230A variant (60% and 17% of the population, respectively) (Fig. S1a-e). To establish CDA as a chemical tool to study the activation mechanism of STING, we first asked whether CDA is a direct or indirect activator of human STING. Similarly to others, we were able to measure human STING binding to cGAMP and CDG, but not to CDA (Fig. S1f). Instead, we turned to the more direct method of protein crystallography. We obtained high-resolution crystal structures of CDA in complex with WT human STING and the 230A variant at resolutions of 2.6 Å and a 2.2 Å, respectively. These structures unequivocally demonstrated that CDA is a direct binder of WT human STING and the 230A variant. We also obtained a structure of cGAMP in complex with the 230A variant at a 1.9 Å resolution (Fig. 1a). In all three ligand-bound human STING structures, we observed the closed STING dimer angle, which differs from the open dimer observed in the apo STING crystal structure (PDB: 4F5W) (Fig. 1b). We quantified the cGAMP- and CDA-induced conformational change by measuring the distance between the tips of the α2 alpha helices (AA 185) of each monomer within the STING dimer in all human STING crystal structures to date (*43-49*). The distances between the two monomers fall into two distinct narrow ranges: 47-54Å (open) and 34-35Å (closed) (Fig. 1c). To eliminate the possibility that the closed dimer is an artifact of crystal packing, we turned to solution phase measurements using small angle X-ray scattering (SAXS) from a synchrotron source. Both cGAMP and CDA binding reduced the radius of gyration (Rg) of STING variants by ∼1.5 Å (Fig. 1d). The decrease in Rg is in agreement with the 1.5Å theoretical reduction in Rg calculated from the crystal structures of apo WT STING and cGAMP-bound WT STING. These results confirm that cGAMP induces closing of the STING dimer in solution, and that CDA induces a similar conformational change in both WT and 230A human STING.

**Fig. 1.**
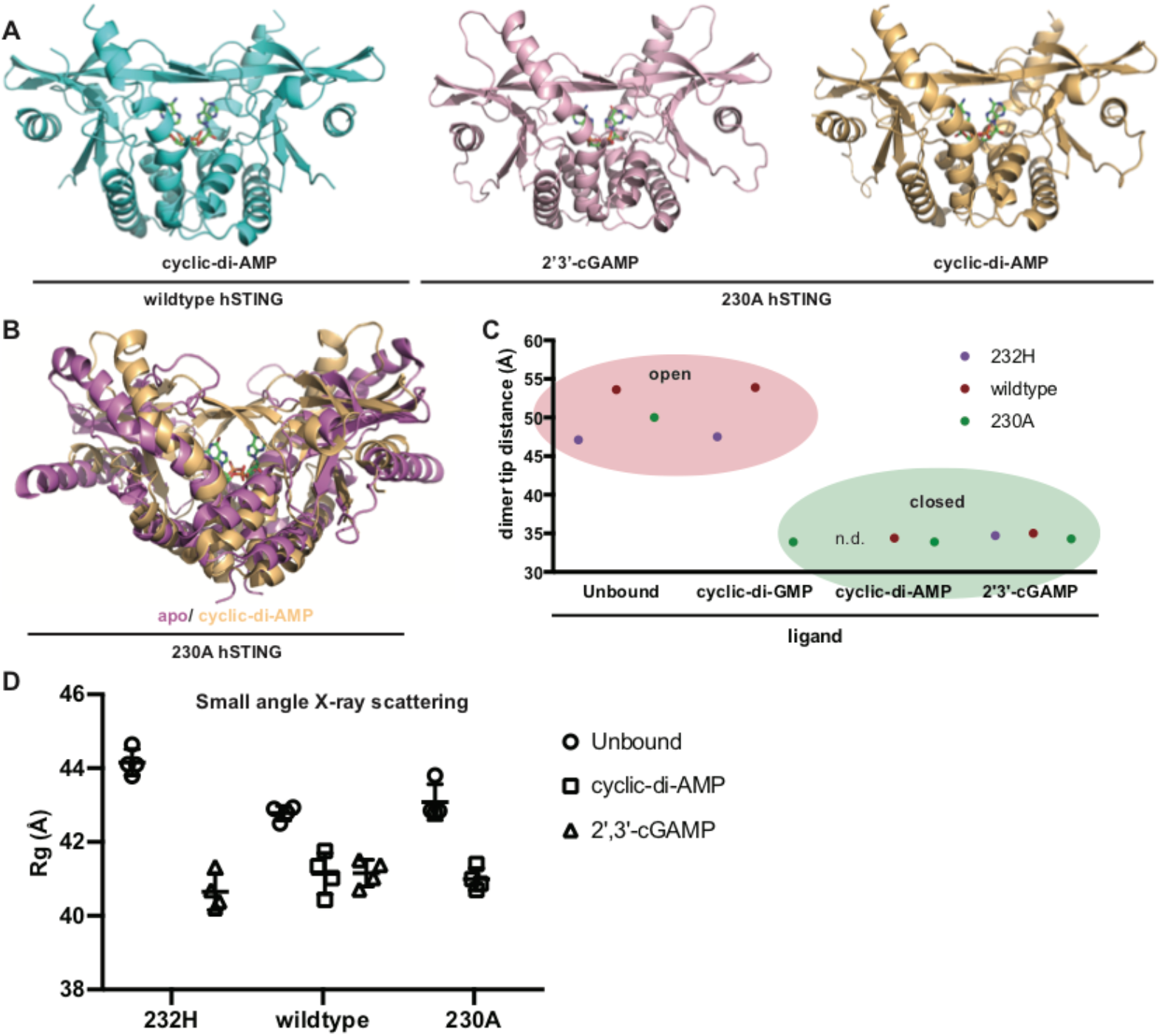
Human STING forms closed dimer angle when bound to cGAMP and CDA. (**A**) Cartoon representations of crystal structures containing the cytosolic domain of human STING variants with indicated ligands (**B**) Overlay of previously solved apo 230A-STING (PDB: 4F5E) with 230A STING:CDA complex. (**C**) Graphical representation of dimer tip distances of all previously solved human STING crystal structures. Distances were measured in angstroms between AA185 of both monomers within a STING dimer. PDB accession numbers of previously solved measured structures are as follows: 4EMU (232H-apo), 4F5W (WT-apo), 4F5E (230A-apo), 4EMT (232H-CDG), 4F5Y (WT-CDG), 4F5D (230A-CDG), 4LOH (232H-cGAMP), 4KSY (WT-cGAMP). (**D**) Small angle X-ray scattering measurements of purified cytosolic human STING variants with or without indicated ligands. Protein was used at a concentration of 1mg/mL with 100 μM ligand. Error bars depict standard deviation from 6 independent replicates.

### STING forms ligand depended polymer on the endoplasmic reticulim

It was previously observed that ligand binding induces human STING aggregation in cells (*12*), but the nature of the STING aggregates and how they lead to activation is not known. We first verified this finding in cells using blue native gel electrophoresis (Fig. 2a). We then determined whether the purified cytosolic domain of STING also aggregates upon ligand binding. Indeed, addition of cGAMP, and to some extent CDA, caused purified STING to shift to higher molecular weights in solution (Fig. 2b). We then examined the crystal lattice of our ligand-bound STING complexes to determine if these ligand-induced STING aggregates formed any ordered structure. Indeed, both CDA- and cGAMP-bound STING formed nearly identical linear polymers in the crystal lattice with their N-termini, which connect to the transmembrane domain, all on the same plane (Fig. 2c, Fig. S2a). In fact, all published human STING crystal structures in the ligand-bound closed conformation form this ordered polymer (Fig. S2b). In contrast, apo STING dimers are stacked top-to-top and bottom-to-bottom in the crystal lattice, a configuration that is geometrically impossible on a membrane (Fig. 2d). In our polymer structure of the cGAMP: G230A STING complex, the homodimer’s total surface area is 29,600 Å^2^ with 8,640 Å^2^ buried in the polymer interface. At this interface, Asp301 from one STING dimer is positioned in between Arg281 and Arg284 from the neighboring dimer, and can form a salt bridge with either residue depending on its orientation within a specific structure. Notably, the salt bridge is formed with Arg284 in the 230A structures and with Arg281 in the WT structures and polymer structure and salt bridge interaction is formed independently from the space group of the crystal lattice. However, when we mutated D301 to alanine, STING was still fully functional, indicating that this interaction is not required for STING function (Fig. 2e). The polymerization interface is vast and it is feasible that many interactions play a role in its stability. We also sought to determine the subcellular location of STING polymerization. It has been shown that STING traffics from the endoplasmic reticulum (ER) to the golgi upon activation, and that disrupting the trafficking with brefeldin A blocks STING signaling (*15).* We found that retaining STING on the ER did not block ligand dependent polymerization (Fig. 2f), indicating that the polymerization event takes place on the ER before trafficking.

**Fig. 2.**
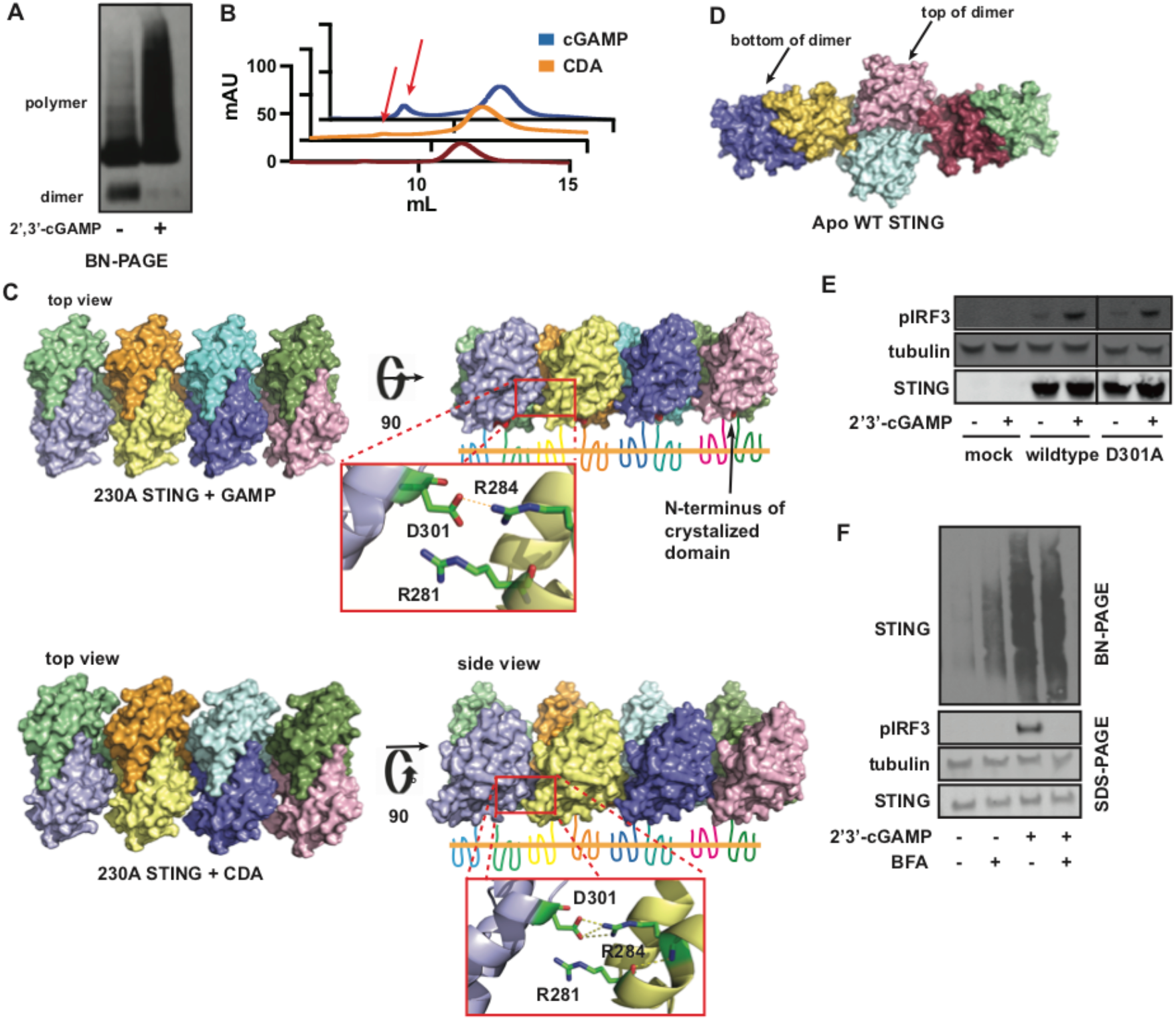
STING forms ligand dependent polymer upon CTT release. (**A**) Blue native PAGE of HEK 293 cells with or without 2 hours stimulation with 100 μM cGAMP. (**B**) Size-exclusion chromatography of 10 μM purified wildtype cytosolic STING incubated at room temperature overnight with and without addition of 500 μM 2’3’-cGAMP or CDA. STING aggregate peaks are indicated with red arrows. (**C**) Structure of crystal packing of 230A STING in complex with cGAMP or CDA. Inlets show alternate salt bridge formation between Asp301 and Arg281 or Arg284. **(D)** Crystal packing of apo (PDB: 4F5W). (**E**) Western blot of HEK 293T cells transfected with indicated plasmids and stimulated for 2 hours with 100 μM cGAMP. (**F**) Western blot and BN-PAGE of U937 cells with or without 1 hour pre-treatment with 40 μM Brefeldin A (BFA) and subsequent 2 hour treatment with 100 μM cGAMP.

### SAVI mutant R284S STING is constitutively polymerized and cannot sequester STING CTT

These interactions drew our attention to three recently reported SAVI-causing STING mutants, C206Y, R281Q, and R284S (*30, 32*), that all reside in the polymerization interface of our crystal structure (Fig. 3a). Another three SAVI-causing STING mutations (V147L, N154S, V155M) reported in an earlier study (*29*) are not near the polymer interface. We, therefore, hypothesized that mutants found in the polymer interface cause constitutive STING polymerization. Indeed, we found that the R284S STING mutant forms constitutive polymers (Fig. 3b). It was previously shown that the STING cytosolic domain binds and sequesters its CTT, but releases the CTT upon CDG binding (*49*). It was hypothesized that freed CTT facilitates STING aggregation. However, since our polymer structures do not involve the CTT, we hypothesized that the CTT binds to and protects the polymer interface in inactive STING. Indeed, when we expressed STING without the CTT (AA 1–343, ΔCTT) it formed a constitutive polymer without ligand activation (Fig. 3c). We then co-immunoprecipitated HA-tagged STINC CTT (CTT-HA, AA 344–379) with WT (ΔCTT-WT STING) or SAVI mutants (ΔCTT-R284S STING and ΔCTT-V147L STING) (Fig. 3d). While ΔCTT-WT STING and ΔCTT-V147L STING co-immunoprecipitated with the CTT, ΔCTT-R284S STING did not (Fig. 3e), suggesting that R284S is unable to bind the CTT, making its polymerization interface constitutively available. Further, addition of cGAMP abolished the interaction of ΔCTT-WT STING and ΔCTT-V147L STING with the CTT (Fig. 3f), indicating that ligand binding triggers CTT release. Together, our results suggest that CTT sequestration in inactive STING prevents constitutive STING polymerization.

**Fig. 3.**
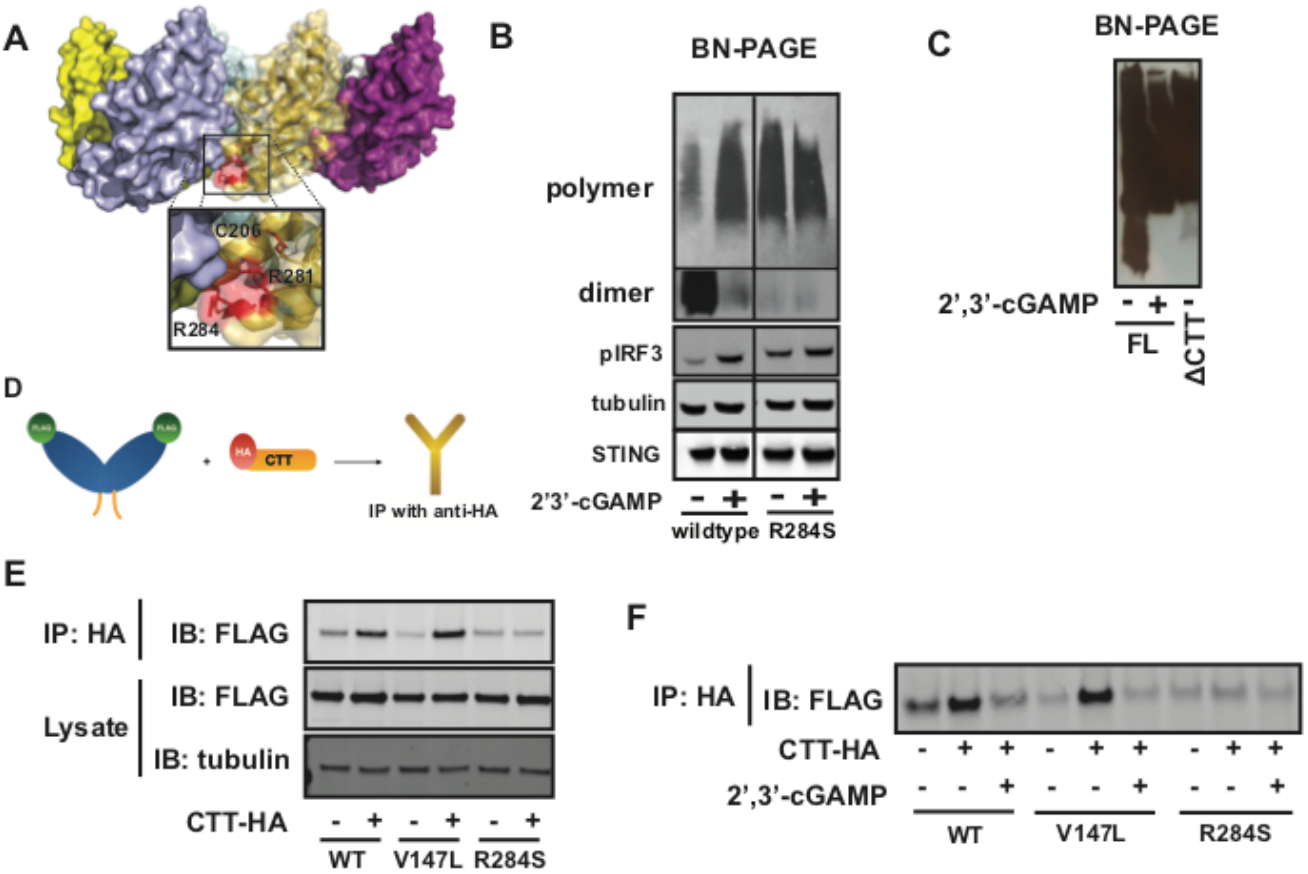
SAVI mutant R284S STING is constitutively polymerized and cannot sequester STING CTT. (**A**) The location of three SAVI-causing STING mutants that reside in the observed polymer interface. (**B**) Blue native PAGE of HEK 293T cells transfected with indicated plasmid and stimulated 2 hours with 100 μM cGAMP. The left and right panels are the same exposure of the same gel with unrelated data omitted from the center, while the top and bottom panels are different exposures of the same gel to observe both the polymer and dimer structures. This is necessary due to a clear antibody preference for polymerized STING. (**C**) Blue native PAGE of HEK 293T cells transfected with equi-molar amounts of STING plasmid, either full length (AA 1-379) or ΔCTT (AA 1-343) and stimulated 2 hours with 100 μM cGAMP. (**D**) Diagram of immunoprecipitation experiment to measure STING cytosolic domain interactions with the STING CTT. (**E**) Immunoprecipitation (IP) of STING CTT followed by immunoblotting (IB) in HEK 293T cells transfected with indicated plasmids. (**F)** The same samples as in (E) with 100 μM cGAMP added to lysates.

### STING polymers are disulfide stabilized

It has been previously observed that STING aggregates do not form in the presence of dithiothreitol (DTT) (*50*), though this observation has not been further explored. We also observed disruption of STING polymers in reducing conditions (Fig. 4a), suggesting that the STING polymer is stabilized by disulfide bonds. We observed a ligand dependent disulfide linkage between the cytosolic domain of two STING molecules using non-reducing PAGE (Fig. 4b). There are five cysteines in the cytosolic domain of STING, four of which (C206, C257, C292, and C309) are buried inside well-defined regions in our structures and are not engaged in disulfide bonds (Fig. S3a). In contrast, C148 resides at the linker region that connects the ligand-binding domain to the transmembrane domain, a flexible 15 amino acid stretch that is not visible in the electron density maps in any of the available STING structures. While we observed cGAMP-induced disulfide bond formation in cells expressing the cytosolic domain of WT STING, we did not when C148 was mutated to alanine (STING-C148A) (Fig. 4c). When purified, the STING-C148A mutant folds correctly and binds cGAMP with only slightly weaker affinity than wildtype (Fig. S3b, c). However, 293T cells transfected with full-length STING-C148A did not respond to cGAMP treatment while those transfected with WT STING did (Fig. 4d). Together, our results support a model in which C148 residues from neighboring STING dimers crosslink and stabilize STING polymers with a well-defined tertiary structure, which is necessary for STING activation (Fig. 4e).

**Fig. 4.**
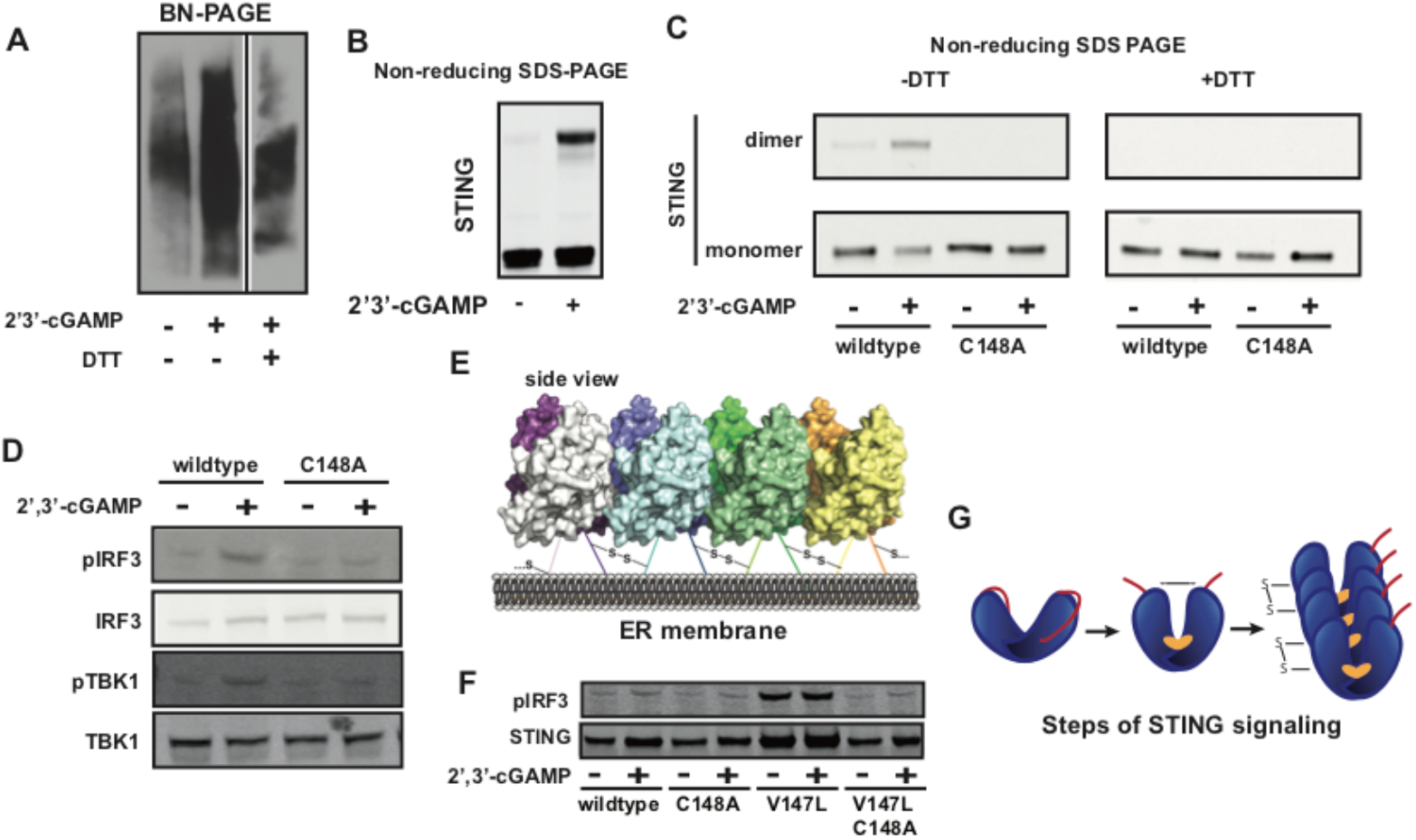
Human STING polymer is disulfide stabilized. (**A**) Blue native PAGE of HEK 293 cells with or without 2 hours stimulation with 100 μM cGAMP and 1 μM DTT added to cells post lysis and solubilization. (**B**) Non-reducing SDS-PAGE of HEK 293T cells transfected with WT cytosolic STING and stimulated for 2 hours with 100 μM cGAMP. (**C)** Non-reducing SDS-PAGE blot of HEK 293T cells transfected with either cytosolic WT or C148A pcDNA3 plasmids and stimulated 2 hours with 100 μM cGAMP. (**D**) Western blot of HEK 293T cells transfected with full-length WT or C148A STING in pcDNA3 plasmids and stimulated 2 hours with 100 μM cGAMP. (**E**) Model of disulfide stabilized STING polymers on the endoplasmic reticulum membrane. **(F)** Western blot of HEK 293T cells transfected with indicated plasmids and stimulated for 2 hours with 100 μM cGAMP. (**G**) Diagram depicting the steps of STING activation by cGAMP. In the final step CTT on only displayed on the right and disulfide bonds only on the left for visual clarity. In reality, both sides of the polymer would have a CTT array and be disulfide linked.

Interestingly, the three SAVI-causing STING mutations that reside outside of the polymer interface (V147L, N154S, V155M) flank C148. While cells expressing V147L-STING exhibited high basal levels of phosphorylated IRF-3, the V147L/C148A-STING double mutant is not constitutively active, nor can it be activated by cGAMP (Fig. 4f). The fact that hotspots for STING disease mutations colocalize with either the polymer interface or C148 further supports that these are key structural regions for STING activation. Together, our results support a model in which cGAMP binding to STING induces a conformational change leading to the release of the CTT, which exposes the polymer interface and, thus, allows disulfide-linked polymer formation (Fig. 4g).

### CDG activates STING cooperatively and is a partial inhibitor of cGAMP signaling

We then asked whether closing of human STING dimer is the conformational change required for STING activation. We turned to CDG as another tool ligand of human STING. Crystal structures of CDG in complex with WT (PDB: 4F5Y), 230A (PDB: 4F5D), or 232H (PDB: 4EMT) STING alleles were the first solved STING crystal structures. In these structures, CDG does not induce conformational changes in WT and 232H STING alleles in reference to the apo STING, but induces dimer closing in the 230A allele. Again, we sought to validate these crystallographic data using SAXS. In solution, CDG binding induced a 1.5Å decrease in Rg in the 230A variant, similar to what we observed with cGAMP binding. Interestingly, addition of CDG to WT STING also decreased the Rg, although to a lesser extent (Fig. 5a). This is in contrast to the crystal structures where the CDG bound and apo WT STING crystal structures overlay completely. We hypothesize that apo STING dimers are more flexible than previously assumed and, hence, have a slightly larger Rg in solution than CDG-bound STING, which would presumably have a more ridged dimer angle. The constraints of the crystal lattice may have stabilized one specific conformation of the apo STING dimer. Given that CDG binding to STING does not cause closing of the dimer, we asked whether CDG is able to activate STING signaling. When we electroporated CDG into primary human lymphocytes harboring wildtype STING, it activated IRF-3 phosphorylation with an EC_50_ of 8 µM (Fig. 5b, c). The EC_50_ is approximately 200-fold weaker than that of cGAMP (Fig. 5d, e), which reflects its 200-fold weaker binding affinity. CDA, which also induces STING dimer closing, activated primary human lymphocytes with an EC_50_ of ∼400 nM (Fig. S4a). Since cGAMP and CDA display lower EC_50_s than CDG, dimer closing likely only contributes to high ligand affinities and is not required for STING activation.

**Fig. 5.**
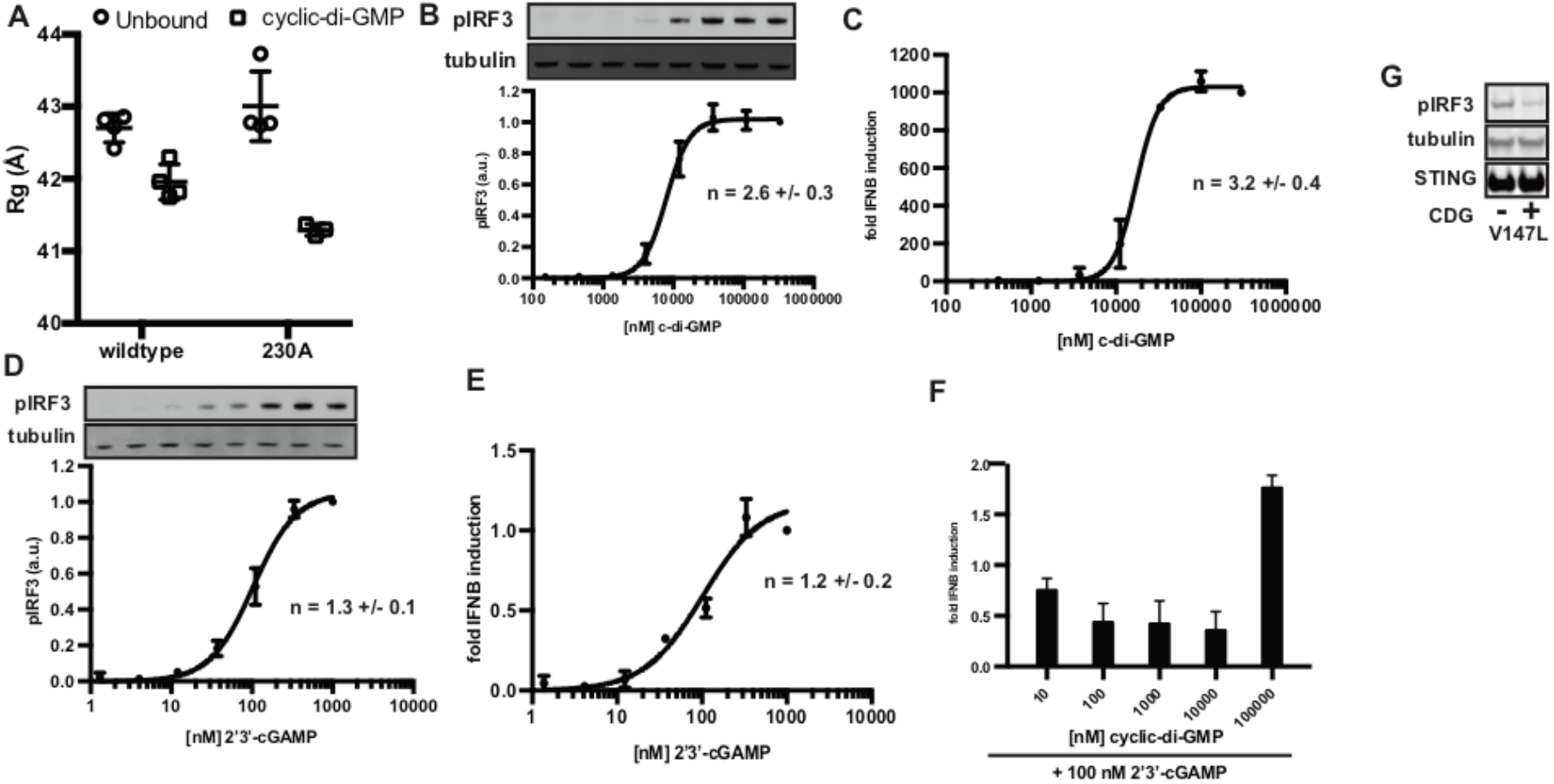
Cyclic-di-GMP activates STING cooperatively and is a partial inhibitor of cGAMP signaling. (**A**) SAXS analysis of STING variants with or without 200 μM CDG. (**B**) Measurement of phosphorylation of IRF3 at various concentrations of CDG electroporated into primary human lymphocytes via western blotting. One representative blot is shown and error bars are generated from the s.d. of three independent replicates. Hill coefficient (n) is measured from fitting the data to a hill equation. (**C**) The same samples as (B) measured by RT-qPCR for *IFNB* mRNA levels. (**D**) and (**E**) are similar to (B) and (C) but with cGAMP treatment instead of CDG. (**F**) *IFNB* mRNA levels measured by RT-qPCR in primary human lymphocytes electroporated with 100 nM cGAMP and indicated concentrations of CDG.

Interestingly, the response curve for CDG is steeper than that for cGAMP, and when fitted with a hill equation, the coefficient for CDG activation is 2.6 ± 0.4 as opposed to a value of 1.3 ± 0.3 for cGAMP, suggesting that CDG signals in a more cooperative manner. Knowing that CDG binding to STING has also been shown to promote STING aggregation (*49*), we predicted that CDG activates STING by inducing polymerization of open STING dimers. Cooperativity is not observed in our in vitro binding assay (Fig. S1f), suggesting that it most likely originates from the polymerization step. Because STING overexpression also leads to ligand-independent activation (Fig. S4b), we hypothesized that inactive STING molecules have affinities towards each other, but due to their flexible dimer angles, would not polymerize unless overexpressed at a high enough concentration on the ER membrane. However, it is possible that CDG-bound STING could polymerize with apo STING, increasing the dimer rigidity of apo STING and therefore increasing its affinity for CDG. This would lead to the observed cooperativity. On the contrary, cGAMP bound STING likely has less affinity for apo STING due to their greater conformational differences and does not signal cooperatively (Fig. S4c). Likewise, due to dimer angle mismatch, CDG-bound STING may not be able to polymerize with cGAMP-bound STING. CDG could therefore act as a competitive inhibitor of cGAMP signaling. Indeed, at a concentration 10-fold below its EC_50_ as a STING agonist, but similar to that of its K_d_, CDG inhibited cGAMP activation of STING by 50% (Fig. 4f). To determine whether CDG inhibition of STING would be relevant in a disease setting, we tested CDG’s ability to inhibit basal activity of SAVI mutant STING-V147L and found that CDG was able to do so by about 40% (Fig. 5g).

## Discussion

In summary, our study yielded a few paradigm shifting discoveries. First, it was thought that closing of the dimer was the key mechanism in human STING activation. Our data, however, demonstrate that the ability of a ligand to induce human STING dimer closing is required for its high potency, but not for its ability to activate STING. Second, we offer structures of the STING polymer, which are different from the previously proposed model (*49*) in that the CTT is not required for polymerization, but rather protects the polymer interface and prevents autoactivation. We also show that the polymerization event occurs at the endoplasmic reticulum, before STING traffics to the golgi. This indicates that the palmitoylation of STING, which occurs at the golgi (*51*), is not required for STING polymerization, but instead serves some other function. In addition, we reported that STING forms a ligand-dependent disulfide bond. Though it is surprising that a disulfide bond could form in the reducing environment of the cytosol, it is very likely that ligand binding induces a conformational change in the transmembrane domain of STING, which either moves C148 into the membrane, or sequesters it between the polymer and the membrane and therefore protects it from reducing agents.

Our model of STING polymerization predicts irreversible STING activation by cGAMP with a high threshold. First, STING could not polymerize unless a certain ratio of STING is occupied by cGAMP. Indeed, the EC_50_ value of cGAMP is 10-fold higher than its K_d_. Second, we observed that the array of CTTs presented on polymerized STING is a better scaffold for IRF3 dimerization than CTTs on unpolymerized STING, since one IRF3 dimer perfectly bridges two polymerized STING dimers (Fig. S4d). Through multivalent interactions, few TBK1 molecules could readily phosphorylate all IRF3 dimers displayed on a STING polymer. Supporting this hypothesis, STING puncta formation and IRF3 nuclear translocation in response to cGAMP both exhibit an all-or-none behavior (*51, 52*). STING’s high threshold of activation is drastically different from transport enzymes and hydrolases which often operate at much lower concentrations than their K_m_ values. However, like STING, cGAS is only activated at enzyme and substrate concentrations much higher than its K_m_ (*53*), suggesting that it is advantageous to have a high threshold for this anti-viral and anti-cancer pathway. Additionally, polymerization as an activating step is also observed in similar innate immune pathways, such as dsRNA receptor MDA5 and adaptor MAVS (*54, 55*). The existence of a threshold of activation could be a central mechanism to distinguish between foreign dsDNA at high acute concentrations and basal levels of self dsDNA. Interestingly, the high threshold of cGAS activation is achieved through a membraneless liquid droplet partitioning mechanism, while in STING is achieved through polymerization on the membrane.

On the translational side, because CDG is able to inhibit cGAMP-induced STING signaling due to its formation of an alternate STING conformation, we predict that any molecule that induces a different STING conformation than cGAMP-bound STING could act as a STING inhibitor. Potent human STING inhibitors that function by preventing its palmitoylation have been developed (*56*). However, we provide a new therapeutic strategy that would prevent low levels of constitutive STING activation in autoimmune and inflammatory diseases, but could be overcome by higher levels of endogenous cGAMP produced during viral infection or cancer invasion. We predict this strategy would be less immunosuppressive, which is crucial for life-long treatment of many autoimmune syndromes.

## Acknowledgments

We thank M. Deller for help initiating SAXS experiments, and all members of the Li lab for helpful feedback and discussions. We thank the staff members of the Macromolecular Crystallography Group at SSRL for excellent beam line support.

## Funding

S.L.E. was supported by NIH T32 Cell and Molecular Biology Training grant 5 T32 GM007276 and the Blavatnik Fellowship. L.L. thanks Ono Pharma Foundation for supporting this research. Use of the Stanford Synchrotron Radiation Lightsource, SLAC National Accelerator Laboratory, is supported by the U.S. Department of Energy, Office of Science, Office of Basic Energy Sciences under Contract No. DE-AC02-76SF00515. The SSRL Structural Molecular Biology Program is supported by the DOE Office of Biological and Environmental Research, and by the National Institutes of Health, National Institute of General Medical Sciences (including P41GM103393). The contents of this publication are solely the responsibility of the authors and do not necessarily represent the official views of NIGMS or NIH.

## Author contributions

S.L.E. and L.L. designed the study. S.L.E, D.F., and T.M.W. carried out experiments. S.L.E. and L.L. wrote the manuscript. All authors discussed findings and commented on the manuscript.

## Competing interests

Authors declare no competing interests.

## Data and materials availability

Accession numbers for crystallography structures in the PDB are: WT STING:CDA **6CFF**, 230A STING:CDA **6CY7**, 230A STING:cGAMP **6DNK**

## Methods

### Reagents, antibodies, and cell culture

2’3’-cGAMP, c-di-GMP, and c-di-AMP were purchased from Invivogen. The following monoclonal antibodies for western blotting were purchased from Cell Signaling Technologies: rabbit anti-phospho-IRF3 (4D4G, 1:1000), rabbit anti-phospho-TBK1/NAK (D52C2, 1:1000), rabbit anti-IRF3 (D83B9, 1:1000), rabbit anti-TBK1/NAK (D1B4, 1:1000), rabbit anti-STING (D2P2F, 1:1000), and mouse anti-tubulin (DM1A, 1:2000). Secondary antibodies were purchased from LI-COR Biosciences: Goat anti-mouse IgG (1:15000) and goat anti-rabbit IgG (1:15000). HEK 293 and U937 cells were purchased from ATCC. Human peripheral blood mononuclear cells (PBMCs) were isolated from whole blood using standard Ficoll procedures. HEK 293 cells were cultured in DMEM (Corning Cellgro) supplemented with 10% FBS (Atlanta Biologics) (v/v) and 100 U/mL penicillin-streptomycin (ThermoFisher). U937 cells were cultured in RPMI (Corning Cellgro) supplemented with 10% heat-inactivated FBS (Atlanta Biologics) (v/v) and 100 U/mL penicillin-streptomycin (ThermoFisher). PBMCs were cultured in RPMI (Corning Cellgro) supplemented with 2% human AB serum (Corning) and 100 U/mL penicillin-streptomycin (ThermoFisher).

### Cell Stimulation and qPCR

HEK 293 cells were plated in 6 well tissue cultured treated plates at 300,000 cells/well. After 24 hours, indicated ligands were added to media. For Brefeldin A treatments, cells were pre-treated for 1 hour pre-stimulation. After 2 hours, cells were lysed directly on plate in 1x Lameli sample buffer (LSB) and phosphorylation of IRF3 was measured through western blotting. U937 cells and PBMCs were electroporated with indicated concentrations of ligands using a Lonza nucleofection kit and nucleofector. After two hours cells were lysed in 1x LSB and analyzed through western blotting or after 16 hours RNA was extracted using standard Trizol (ThermoFisher) extraction procedures. cDNA was generated using Maxima Reverse Transcriptase (ThermoFisher), and qPCR was carried out using AccuPower 2x Greenstar qPCR Master Mix.

### Western blotting

Cell lysates were separated on a SDS-polyacrylamide gel (Invitrogen), and transferred to a nitrocellulose membrane using a wet transfer system (BioRad). Primary antibody was added overnight at 4°C, followed by three washes in 1xTBS-0.1% Tween. Secondary antibody was added for 1 hour at room temperature, followed by three additional washes in TBS-T. Blots were imaged in IR using a Li-Cor Odyssey Blot Imager. Bands were quantified using ImageJ.

### Co-immunoprecipitation

HEK 293T cells were plated in a 12-well plate (Corning) at 100,000 cells/ well. After 24 hours cells were transfected with pcDNA3 plasmids containing ΔCTT STING-FLAG variants with or without co-transfection with pcDNA3 CTT-HA plasmids using Fugene 6 Transfection Regent (Promega). After 24 additional hours cells were lysed in mild detergent (10% glycerol 25mM NaCl, 20mM HEPES pH 7.0, 1% DDM, + 1x protease inhibitor (Roche)), lysates were pre-cleared for 20 minutes with magnetic protein A beads (CST), and primary HA antibody was added (1:50) overnight. Lysates were incubated with beads 20 minutes RT and washed 5 times in.1% DDM buffer. Protein was eluted from beads in LSB and boiled 10m at 95oC. Protein was analyzed with western blotting.

### Blue Native-PAGE

Cells were lysed in native lysis buffer (10% glycerol, 25mM NaCl, 20mM HEPES pH 7.0) + 1%DDM and protease inhibitor (Roche) and solubilized by rotating 30m at 4°C. Lysates were spun 10m at 14000 rpm and supernatant was collected. Lysate was added to 4x native sample buffer (Invitrogen), run on a NativePAGE gel (Invitrogen), and transferred to a PVDF membrane using a wet transfer system (BioRad). Western blotting protocol was followed for membrane staining.

### STING cloning, protein purification, and cGAMP binding assay

DNA sequence encoding for cytosolic domain of human STING was PCR amplified from HEK 293 cell cDNA using Phusion High-Fidelity DNA polymerase (Thermo). The PCR product was inserted into SapI and XhoI sites of pTB146 using isothermal assembly. Alternate hSTING variants were generated using site-directed mutagenesis. His-tagged STING CTDs were purified as previously described (*39*). Protein was further purified using size-exclusion chromatography on a superpose 12 column (GE Healthcare) run on an Äkta pure system (GE Healthcare). In solution STING aggregation was measured using this system. Corrected folding was checked using Circular Dichroism as previously described (*57*) and cGAMP binding was verified using the radioactivity based nitrocellulose filter binding assay. 35S-labeled 2′3′-cGAsMP was used as a competitive probe and synthesized as previously described (*17*).

### STING Structure Determination

Human STING cytosolic domain residues 133-379was overexpressed in E. coli. The protein has been purified to homogeneity and concentrated to 10 mg/ml prior to crystallization screening. The screens were designed from available crystal structure determinations, and custom-made screens were made based on a broad range of molecular weight PEGs. In parallel, commercial screens based on this precipitant were also employed. The vapor-diffussion method, employing sitting-drop hSTING co-crystallization (using 1mM CDN) was performed on 96-well microtiter plates at three different temperatures. WT-hSTING:CDA needle-shaped crystals appeared within a week of setting up the crystallization trays at 4 and 12 °C. Multiple hits from many different conditions were observed but, when X-ray exposed, these crystals had weak diffracting power. Some of the conditions were chosen for a second-round fine-screening and, within a month, better quality crystals appeared. A dataset to a minimal Bragg spacing of 2.4 Å was collected from a crystal that grew from a condition composed by 0.2 M Sodium citrate tribasic, 20 % PEG 3350, pH 8.2. The crystal belonged to the tetragonal space group P 4_1_ 2_1_ 2 and contained one polypeptide chain per asymmetry unit. The structure was solved by the molecular replacement method with Phaser (*58*) using the polypeptide chain of hSTING (PDB: 4LOH) as the search model. Presence of extra electron density was evident after structure solution and accounted for half a CDA molecule (the other half is generated via a space group symmetry operation). Of the single independent polypeptide chain, residues 154-335 were unambiguously traced in the electron density maps except for loop regions 184-193 and 317-324 for which very weak electron density prevented defining an accurate model. A similar strategy was adopted to obtain CDA-G230A-hSTING and cGAMP-G230A-hSTING crystal structures. Needle-shaped crystals were grown from conditions consisting of: 0.2 M Ammonium citrate tribasic, 0.1M imidazole, 20 % PEG MME 2000, pH 7.0 and 0.2 M ammonium sulfate, 20% PEG 3350, pH 6.7, respectively. Data was collected to minimal Bragg spacings of 2.2 Å and 1.9 Å, respectively, at SSRL beam line 12-2. Crystals were in the same tetragonal space group with one polypeptide chain per asymmetry unit. Imposed by symmetry, either the GMP and AMP moieties in cGAMP have been modeled interchangeably at 0.5 occupancy in the binding pocket of the cGAMP-G230A-hSTING crystal structure. As with other human STING structures, N-terminal residues up to 154 and the C-terminal tail beyond residue 336 were not visible in the maps and not modeled. To generate accurate bond length, angle, and torsion restraints for the CDA molecule, preliminary refinement rounds using SHELXL (*59*) were performed and the parameters later used in the final steps of refinement with REFMAC (*60*). During refinement manual adjustments on the polypeptide chain were made to account for extra electron density peaks in COOT (*61*). Solvent water molecules were first assigned based on their hydrogen bonding properties to the CDN and its surrounding residues; in later stages of refinement, further water molecules were added automatically. Refinement progressed to convergence and reached an excellent agreement between the model and the experimental data (see Table 1). Crystals were harvested and cryocooled under the LN_2_ stream at the beam line and datasets were collected at SSRL synchrotron (*62*). Data was reduced with XDS (*63*), scaled with SCALA (*64*) and analyzed with different computing modules within the CCP4 suite (*65*). Graphic renderings were prepared with pymol (*66*). Crystals from the CDA-R232H-hSTING variant were also obtained but they diffracted poorly, preventing any structural determination.

**Table 1.**
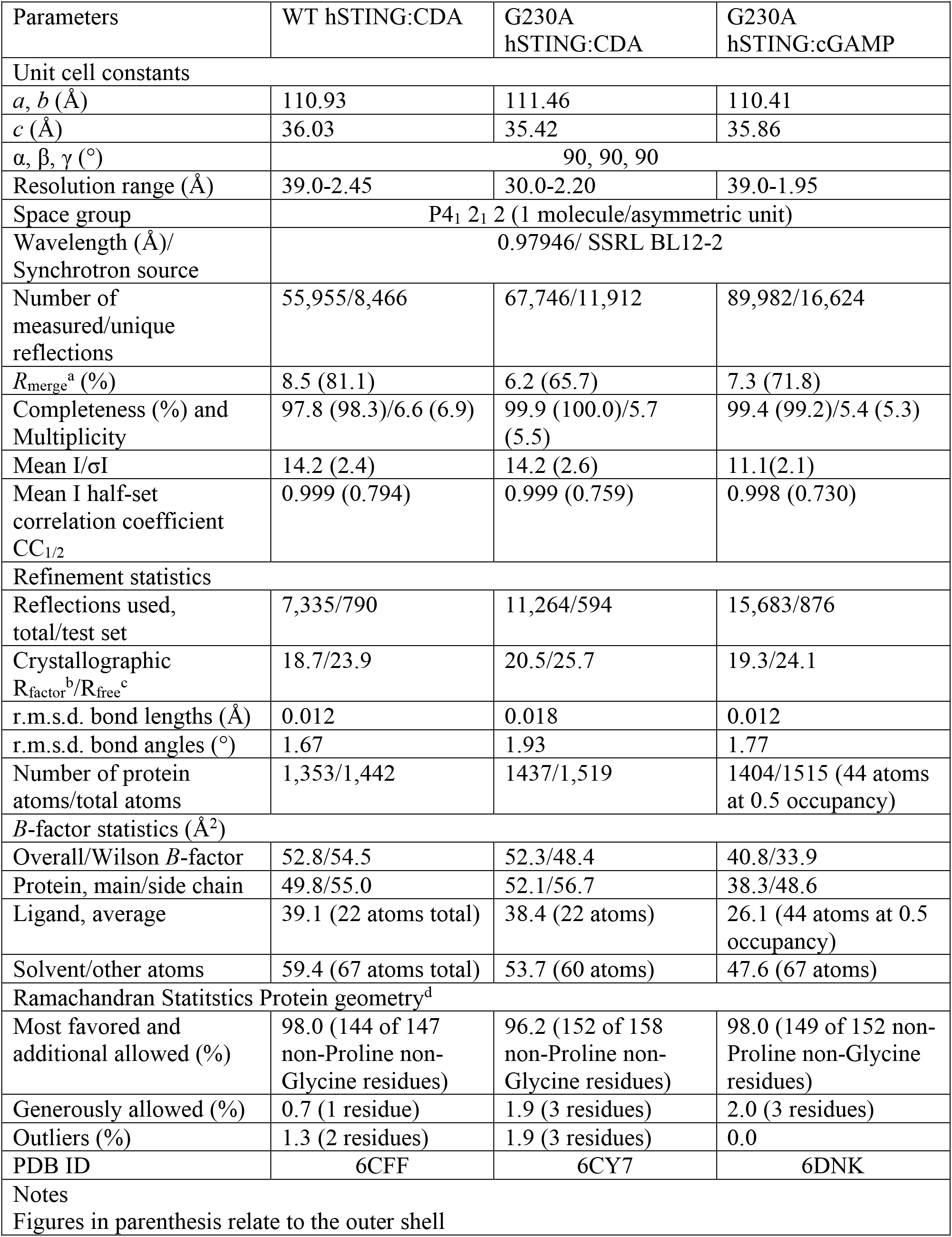
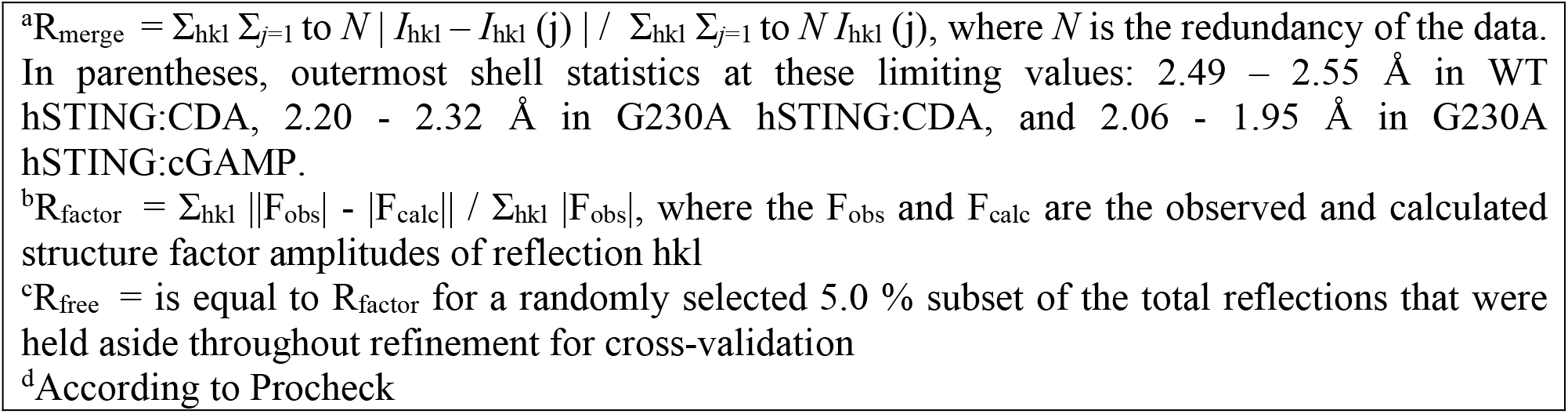
Crystallographic Data Statistics

| **Table 1. Crystallographic Data Statistics** |  |  |  |
| --- | --- | --- | --- |
| Parameters | WT hSTING:CDA | G230A hSTING:CDA | G230A hSTING:cGAMP |
| Unit cell constants |  |  |  |
| $a, b$ (Å) | 110.93 | 111.46 | 110.41 |
| $c$ (Å) | 36.03 | 35.42 | 35.86 |
| $\alpha, \beta, \gamma$ (°) | 90, 90, 90 | | |
| Resolution range (Å) | 39.0-2.45 | 30.0-2.20 | 39.0-1.95 |
| Space group | P4<sub>1</sub> 2<sub>1</sub> 2 (1 molecule/asymmetric unit) |  |  |
| Wavelength (Å)/<br>Synchrotron source | 0.97946/ SSRL BL12-2 |  |  |
| Number of<br>measured/unique<br>reflections | 55,955/8,466 | 67,746/11,912 | 89,982/16,624 |
| $R_{\text{merge}}^a$ (%) | 8.5 (81.1) | 6.2 (65.7) | 7.3 (71.8) |
| Completeness (%) and<br>Multiplicity | 97.8 (98.3)/6.6 (6.9) | 99.9 (100.0)/5.7 (5.5) | 99.4 (99.2)/5.4 (5.3) |
| Mean $I/\sigma I$ | 14.2 (2.4) | 14.2 (2.6) | 11.1(2.1) |
| Mean $I$ half-set<br>correlation coefficient<br>$CC_{1/2}$ | 0.999 (0.794) | 0.999 (0.759) | 0.998 (0.730) |
| Refinement statistics |  |  |  |
| Reflections used,<br>total/test set | 7,335/790 | 11,264/594 | 15,683/876 |
| Crystallographic<br>$R_{\text{factor}}^b/R_{\text{free}}^c$ | 18.7/23.9 | 20.5/25.7 | 19.3/24.1 |
| r.m.s.d. bond lengths (Å) | 0.012 | 0.018 | 0.012 |
| r.m.s.d. bond angles (°) | 1.67 | 1.93 | 1.77 |
| Number of protein<br>atoms/total atoms | 1,353/1,442 | 1437/1,519 | 1404/1515 (44 atoms<br>at 0.5 occupancy) |
| $B$-factor statistics (Å<sup>2</sup>) | | | |
| Overall/Wilson $B$-factor | 52.8/54.5 | 52.3/48.4 | 40.8/33.9 |
| Protein, main/side chain | 49.8/55.0 | 52.1/56.7 | 38.3/48.6 |
| Ligand, average | 39.1 (22 atoms total) | 38.4 (22 atoms) | 26.1 (44 atoms at 0.5<br>occupancy) |
| Solvent/other atoms | 59.4 (67 atoms total) | 53.7 (60 atoms) | 47.6 (67 atoms) |
| Ramachandran Statistics Protein geometry<sup>d</sup> |  |  |  |
| Most favored and<br>additional allowed (%) | 98.0 (144 of 147<br>non-Proline non-Glycine residues) | 96.2 (152 of 158<br>non-Proline non-Glycine residues) | 98.0 (149 of 152 non-Proline non-Glycine<br>residues) |
| Generously allowed (%) | 0.7 (1 residue) | 1.9 (3 residues) | 2.0 (3 residues) |
| Outliers (%) | 1.3 (2 residues) | 1.9 (3 residues) | 0.0 |
| PDB ID | 6CFF | 6CY7 | 6DNK |
| Notes |  |  |  |
| Figures in parenthesis relate to the outer shell |  |  |  |

<sup>a</sup> $R_{\text{merge}} = \sum_{hkl} \sum_{j=1 \text{ to } N} |I_{hkl} - I_{hkl}(j)| / \sum_{hkl} \sum_{j=1 \text{ to } N} I_{hkl}(j)$ , where $N$ is the redundancy of the data. In parentheses, outermost shell statistics at these limiting values: 2.49 – 2.55 Å in WT hSTING:CDA, 2.20 - 2.32 Å in G230A hSTING:CDA, and 2.06 - 1.95 Å in G230A hSTING:cGAMP.
<sup>b</sup> $R_{\text{factor}} = \sum_{hkl} ||F_{\text{obs}}| - |F_{\text{calc}}|| / \sum_{hkl} |F_{\text{obs}}|$ , where the $F_{\text{obs}}$ and $F_{\text{calc}}$ are the observed and calculated structure factor amplitudes of reflection $hkl$
<sup>c</sup> $R_{\text{free}}$ = is equal to $R_{\text{factor}}$ for a randomly selected 5.0 % subset of the total reflections that were held aside throughout refinement for cross-validation
<sup>d</sup>According to Procheck

### Small-angle X-ray scattering

Synchrotron SAXS data were collected at beamline 4-2 (*67)* of the Stanford Synchrotron Radiation Lightsource (SSRL), Menlo Park, CA. The sample to detector distance was set to 1.7m with and X-rays wavelength of λ = 1.127 Å (11keV). Using a Pilatus3 × 1M detector (Dectris Ltd, Switzerland) the setup covered a range of momentum transfer q ≈ 0.006 – 0.51 Å^-1^ where q is the magnitude of the scattering vector defined as q = 4π sinθ /λ, with θ the scattering angle and λ the wavelength of the X-rays. Aliquots of 30 µl of freshly extruded vesicles were loaded onto the automated sample loader (*68)* at the beamline. Consecutive series of sixteen 1s exposures were collected first from the buffer blank followed by the samples. A second buffer blank was measured at the end of each sample series in order to confirm that the sample cell has been cleaned correctly. Solutions were oscillated in a stationary quartz capillary cell during X-ray exposure to increase the exposed sample volume and therefore reduce the radiation dose per protein molecule. The collected data was radially integrated, analyzed for radiation damage and buffer subtracted using the fully automated data reduction pipeline at the beam line. To improve statistics and check for repeatability of the data the measurements were repeated with different sample aliquots 3 to 4 times per sample. As no significant differences were found between the repeat measurements, the different data sets for each sample were averaged.

## Supplemental Figures S1-4 and Table S1

**Fig. S1.**
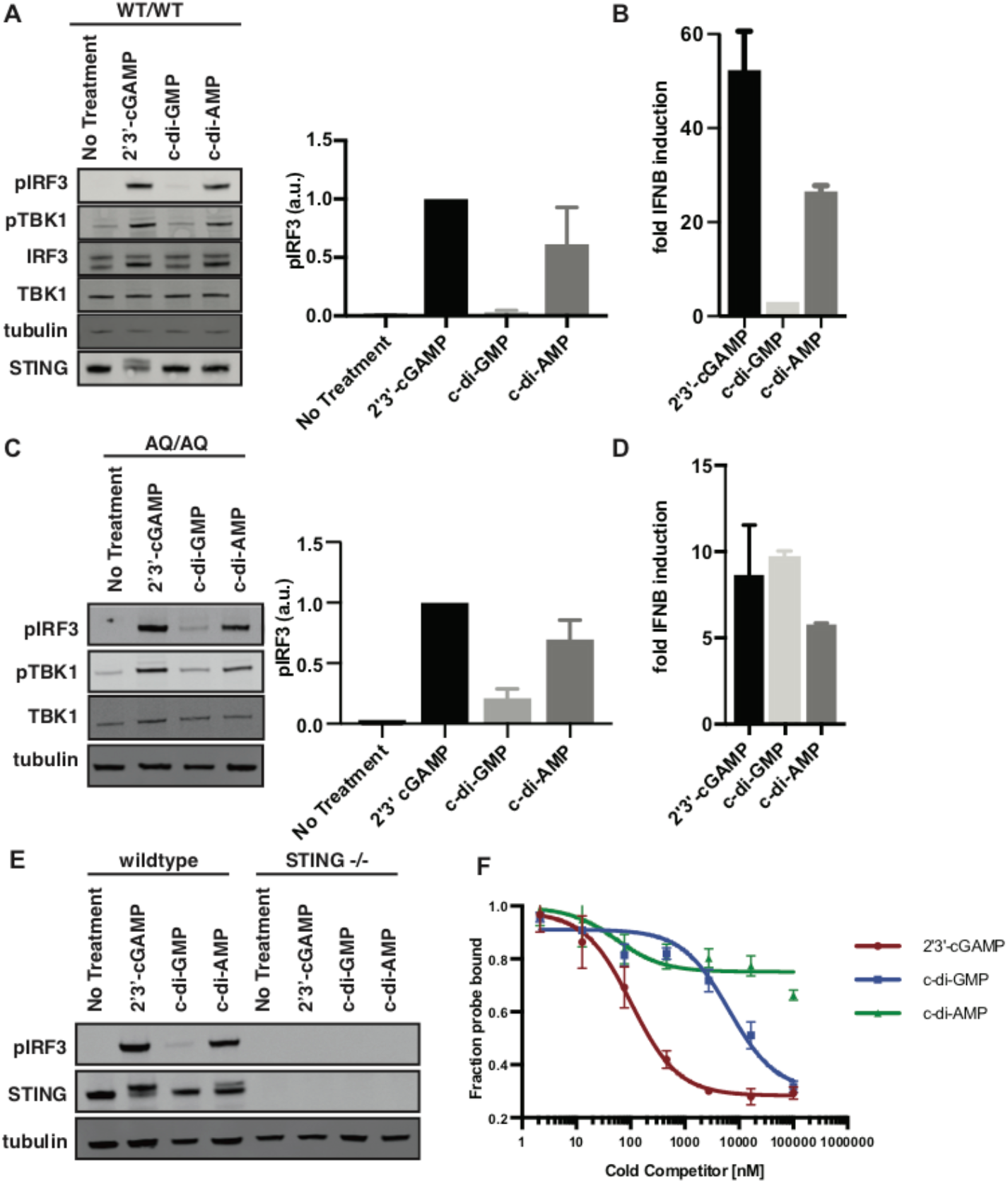
CDA activates WT and AQ STING. (**A**) Western blots of wildtype human primary leukocytes electroporated with 5 μM indicated ligands. Bar graphs indicate s.d. measured from three independent replicates. (**B**) RT-qPCR of *IFNB* mRNA levels in WT primary human leukocytes. (**C**) and (**D**) are similar to (A) and (B) except with primary human leukocytes isolated from individuals homozygous for the 230A 293Q allele. (**E**) Wildtype and STING knockout U937 cells electroporated with 5 μM of indicated ligand. (**F**) Radioactive nitrocellulose filter binding assay of cGAMP, CDG, and CDA competing S35-cGAMP for binding to purified wildtype STING. Error bars are representative of three independent replicates.

**Fig. S2.**
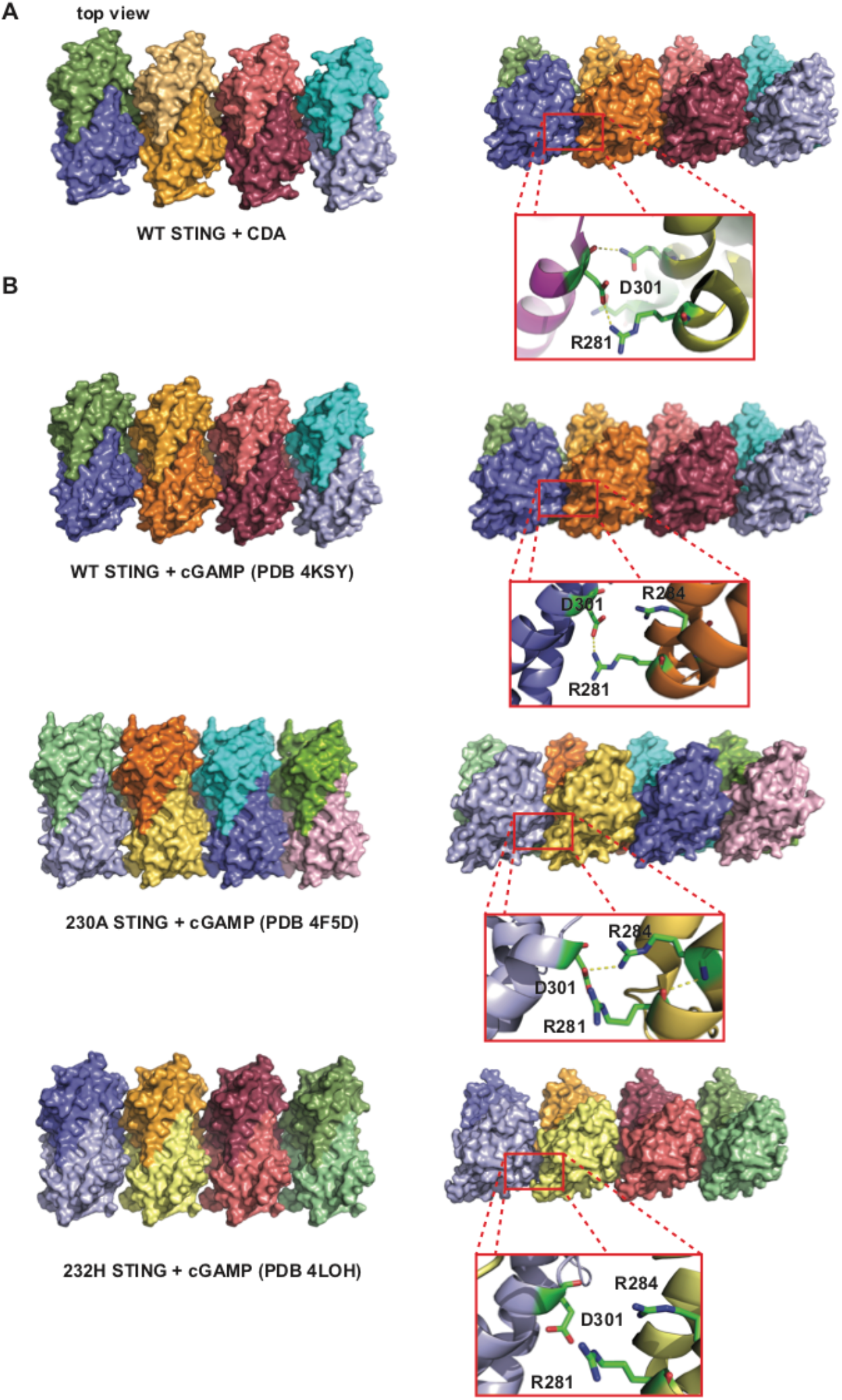
CDA bound WT forms ordered polymers in crystal structures. (**A**) Top view of crystal packing of WT STING with crystalized with CDA. (**B**) Crystal packing of previously published STING structures.

**Fig. S3.**
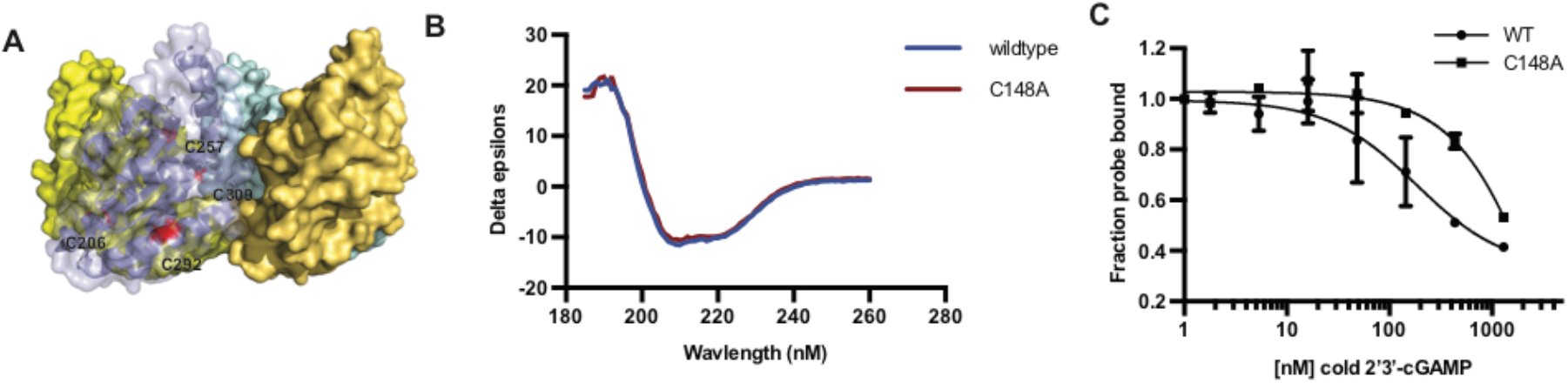
C148A mutant folds correctly and binds cGAMP. (**A**) Location of 4 cysteines visible in cytosolic human STING crystal structures, none of which are engaged in disulfide bonds. (**B**) Circular dichroism of WT and C148A human STING. (**C**) Nitrocellulose filter binding assay of cold 2’3-cGAMP competing S35-cGAMP for STING binding.

**Fig. S4.**
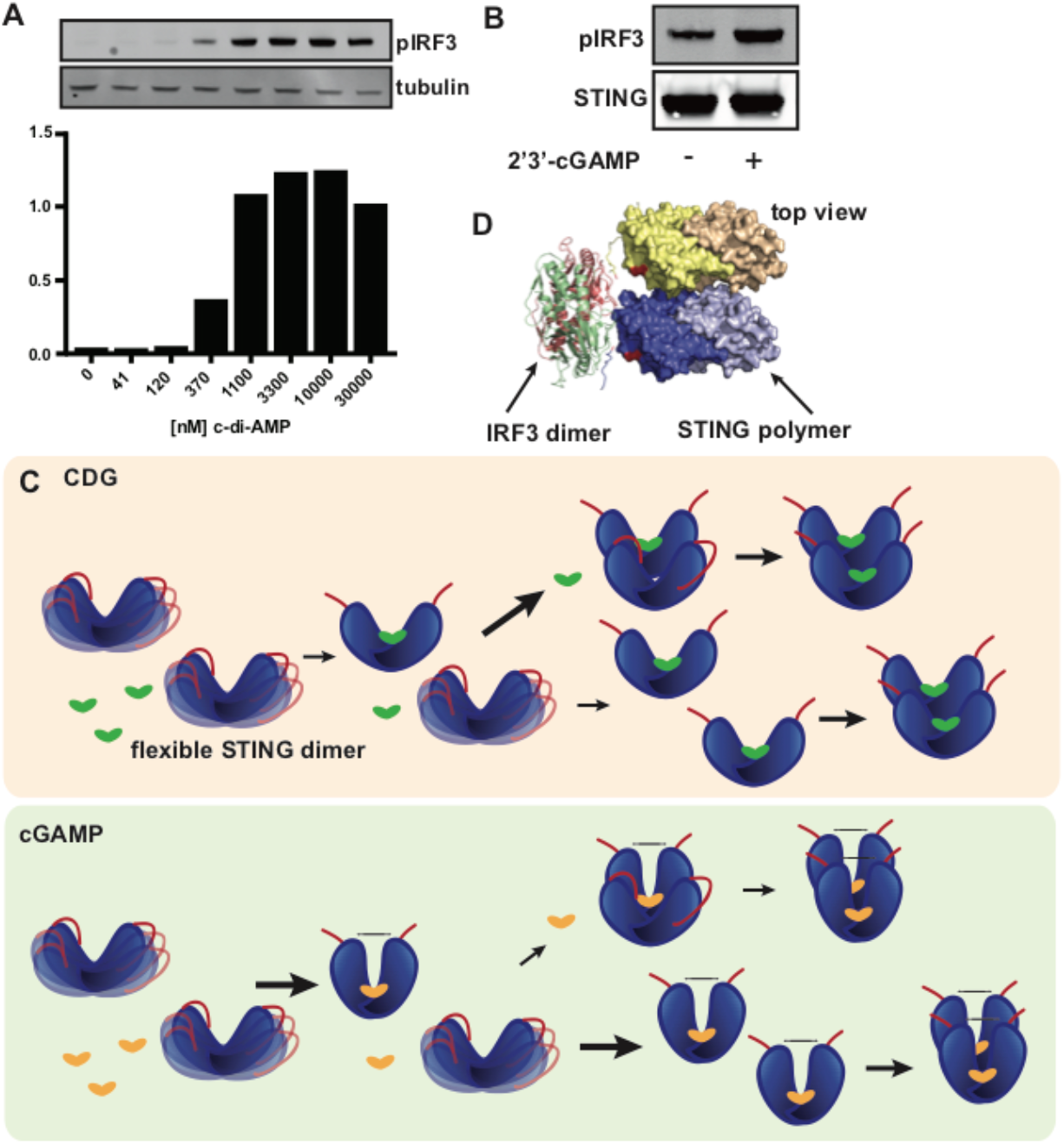
CDG binds STING with less affinity than cGAMP and activates STING cooperatively**. (A)** Measurement of phosphorylation of IRF3 at various concentrations of CDA electroporated into primary human lymphocytes via western blotting. Quantification of western blot is depicted below. **(B)** Western blot of phosphorylation of IRF3 in HEK 293T cells transfected with high levels of WT STING plasmid. **(C)** Diagram of model for cooperative activation of STING by CDG. CDG binds STING in an open conformation that is slightly smaller than the unbound, more flexible, STING. Polymerization of a CDG bound STING to an unbound STING molecule is more favorable than a second CDG binding event, and therefore polymers form with only one bound CDG. Subsequently CDG binding to the polymer are more favorable than the initial CDG binding event, which leads to the observed cooperativity. For cGAMP, the second binding event is more favorable than cGAMP-bound STING polymerizing with unbound STING since 1) the binding affinity of cGAMP to STING is greater, and 2) closed, cGAMP bound STING is more different in conformation to unbound STING than CDG-bound STING to unbound STING. Therefore, cGAMP bound STING polymers form after all the binding events have occurred and therefore no cooperativity is observed. **(D)** Top view of the crystal structure of IRF3 dimer with STING CTT (PDB: 5JEJ) docked onto our polymer structure. The end of our structure and the beginning of the STING CTT co-crystalized with IRF3 are indicated in red.

**Table S1.**

Crystallographic statistics for WT:CDA, 230A:CDA, and 230A:cGAMP structures

